# LoRTE: Detecting transposon-induced genomic variants using low coverage PacBio long read sequences

**DOI:** 10.1101/073551

**Authors:** Eric Disdero, Jonathan Filée

**Affiliations:** Laboratoire Evolution, Génomes, Comportement, Ecologie; CNRS, IRD, Université Paris-Saclay, Gif-sur-Yvette, France

## Abstract

**Motivation:** Population genomic analysis of transposable elements has greatly benefited from recent advances of sequencing technologies. However, the propensity of transposable elements to nest in highly repeated regions of genomes limits the efficiency of bioinformatic tools when short read sequences technology is used.

**Results:** LoRTE is the first tool able to use PacBio long read sequences to identify transposon deletions and insertions between a reference genome and genomes of different strains or populations. Tested against Drosophila melanogaster PacBio datasets, LoRTE appears to be a reliable and broadly applicable tools to study the dynamic and evolutionary impact of transposable elements using low coverage, long read sequences.

**Availability and Implementation:** LoRTE is available at http://www.egce.cnrs-gif.fr/?p=6422. It is written in Python 2.7 and only requires the NCBI BLAST + package. LoRTE can be used on standard computer with limited RAM resources and reasonable running time even with large datasets.

## 1 INTRODUCTION

Transposable elements (TEs), which represent an essential part of eukaryotic and prokaryotic genomes, play important roles in genome size, structure and functions (Fedoroff, 2012; Hua-Van, et al., 2011). TE identification and annotation remains one of the most challenging task in computational genomics but our knowledge of the TE diversity and dynamics among genomes has greatly benefited from the recent advance of sequencing technologies (Lerat, 2010). Specifically, comparison of closely related strains or species using short read sequencing technologies enabled new insights into TE dynamic and their roles in generating structural genomic variation. Two different approaches with their associated computational tools have been developed to achieve this goal. The first approach is based on the direct assembly of the repeated fraction of the reads using highly abundant k-mer: RepARK (Koch, et al., 2014) or Tedna (Zytnicki, et al., 2014) for example. Other tools such as RepeatExplorer (Novak, et al., 2013) or dnaPipeTE (Goubert, et al., 2015) on low-coverage sub-samples of genomic reads to retrieve and specifically assemble the highly repeated elements. All these tools have the advantage to give a good picture of the global TE abundance and diversity. However they do not provide the exact genomic positions of each TE, preventing the identification of the presence/absence of given TE copies between related populations or species. The second approach is implemented in programs that have been specifically developed to detect transposon insertion/deletion between a reference genome and Illumina or 454 short read sequences (Fiston-Lavier, et al., 2015; Kofler, et al., 2012; Rahman, et al., 2015; Zhuang, et al., 2014). The global architecture of these softwares is similar: 1. New insertions are detected by retrieving the reads that do not map on the reference genomes but that align both on a TE consensus sequence and a unique region in the genome. 2. Deletions are detected by identifying reads that span the two flanking sequences of a given TE but do not contain the sequence of the TE copy. Programs like TIDAL also take advantage of the presence of paired end sequences on Illumina reads to identify the deleted locus (Rahman, et al., 2015). This later approach has been extensively tested and benchmarked on diverse *Drosophila* datasets leading to mixed results. Indeed, comparison of respective performance of each program indicated that a very small fraction of the TE insertion/deletion were identified by all programs (Rahman, et al., 2015; Song, et al., 2014). For example, the comparison of TIDAL, TEMP, LnB and CnT on DGRP *Drosophila* strains revealed that only 3% of the calls are predicted in common by the different programs. Thus, a large majority of the predictions are program-specific and PCR validations of the calls lead to substantial levels of false positive (around 40%)(Rahman, et al., 2015). These limitations are mainly due to the fact that TEs tend to insert preferentially in highly repetitive regions. The short length of Illumina reads prevents the precise identification and mapping of these TEs nested in one another. Interestingly, long read sequencing technologies such as those provided by PacBio or MinION technologies are now generating read length that may span the entire length of full transposons and their associated flanking genomic sequences. However, existing programs are not designed to deal with long read sequences and the implementation of new methods is thus required. Here we present LoRTE (Long Read Transposable Element), the first tool for population genomic analyses of TE insertion/deletion between a reference genome and PacBio long read sequences.

## 2. RESULTS AND CONCLUSION

Implementation and dataset details are given in Supplementary Fig S1 and Supplementary Material.

The tool was benchmarked on two *D. melanogaster* PacBio dataset. The first is a synthetic dataset composed of 3 to 30kb PacBio-like reads generated from the reference genome in which we inserted and deleted respectively 100 and 250 TEs. The second is a real biological dataset with *D. melanogaster* PacBio reads coming from the same strain used in the reference genome. We first tested the ability of LoRTE to provide variant calls on a list of 4239 annotated TEs with respect to the read coverage (Fig 1A). For both datasets, LoRTE was able to provide a decision for >99% of the TE locus with a coverage of 9x. Due to the relatively high error rate of the genuine PacBio raw read (around 10%, mainly short insertion/deletion events), synthetic reads performed better at low coverage. Moreover, LoRTE achieved a complete analysis of the data with 10x coverage on a standard computer running at 2.3 GHz in less than 48h, using a maximum of 8 Gb of RAM. This result indicate that a low PacBio read coverage, corresponding to a single single-molecule real-time (SMRT) cell generating 500 to 1000Mb of sequences, is sufficient to make a call on the vast majority of the TE identified in the *D. melanogaster* genome.

**Figure 1:**
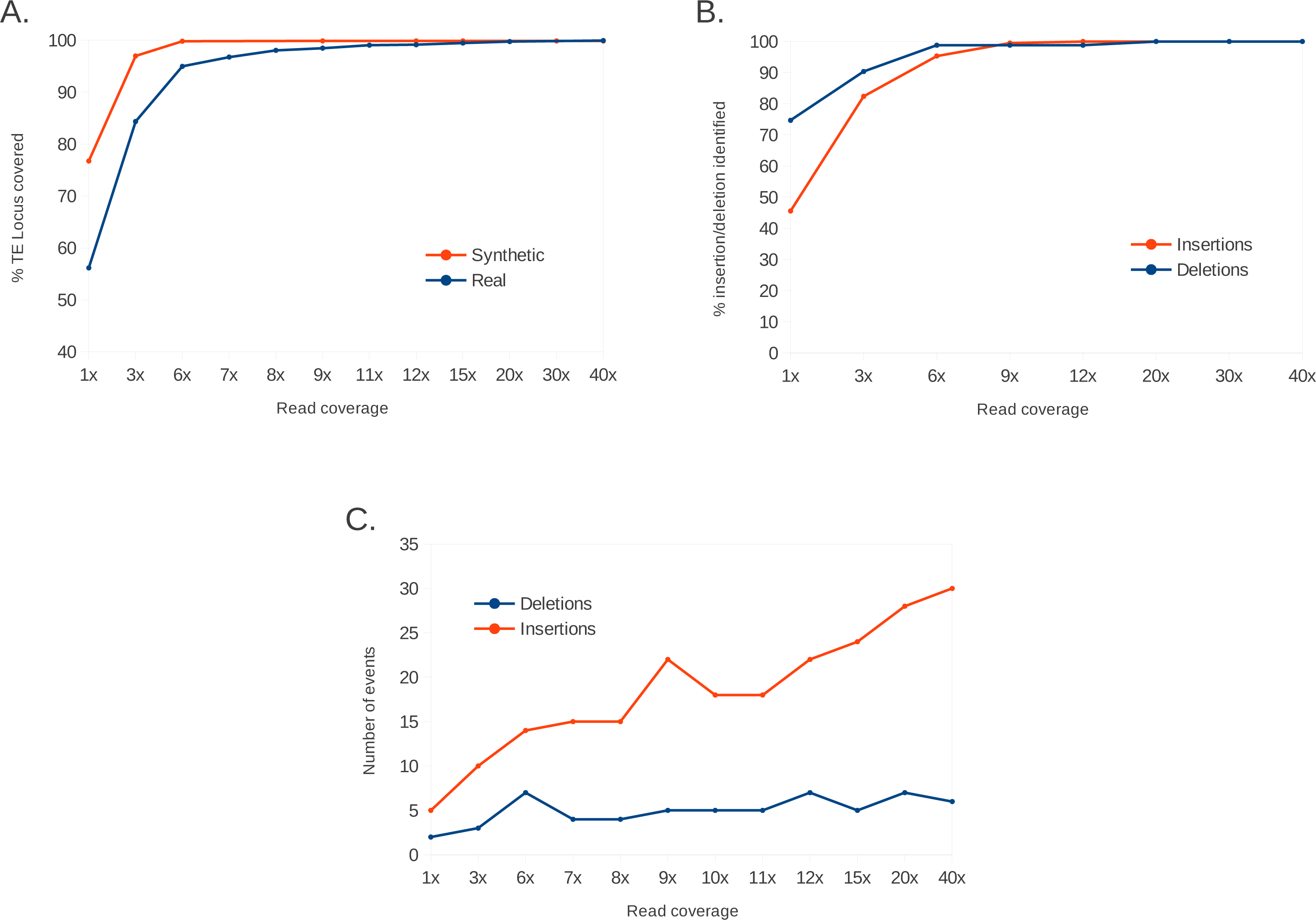
Performance test of LoRTE according to the PacBio read coverage. (A) Percentage of the TEs annotated in the *Drosophila melanogaster* genome that have been recovered by the program. (B) Percentage of the insertion/deletion artificially made in the synthetic reads that have been identified. (C) Numbers of new TE deletion and insertion found in the genuine reads and absent in the reference genome.

We then tested the ability of LoRTE to detect the insertions/deletions made on the synthetic dataset. Fig 1B displays the percentage of insertions/deletions detected by LoRTE with respect to the read coverage. LoRTE detected 98% of the deletions and 100% of the insertion from coverage of 9X and did not generated false positive calls, whatever the coverage. This result strengthens the reliability of LoRTE, even in a context of low coverage PacBio datasets.

We finally analyzed the results obtained by LoRTE on genuine *D. melanogaster* PacBio reads. Fig 1C shows the number of deletion/insertion found in these reads. The number of deletions was relatively constant whatever the read coverage considered. We identifed a maximum of seven deletions corresponding mainly to LTR retrotransposons (two *roo*, two *297*, one *412*), one LINE (*I* element) and one hAT DNA transposon (supplementary Fig S2). All these deletions were present in the 90x genome assembly suggesting that these variants are *bona fide* TE deletions that were not present in the reference genome. Conversely, the number of new TE insertions observed in the PacBio reads increased linearly with the reads coverage’s (Fig 1C). The vast majority of these new insertions are due to *Hobo* elements, a hAT DNA transposon known to have been recently acquired in *D. melanogaster* and subject to a fast and ongoing expansion in the genome (Ragagnin, et al., 2016)(Supplementary Fig S2). Among the 30 new insertions identified using a coverage of 40X, only 12 were validated in the 90X genome assembly. The remaining 18 insertions were absent in the assembly and their calls were supported by only one or a few PacBio reads. These insertions most probably result from somatic insertions at low frequencies but possible false positives could not be ruled out.

Taken together, our results indicate that LoRTE is an efficient and error-free tool to identify structural genomic variants caused by TE insertion or deletion among closely related populations or strains. Here, we demonstrated that LoRTE performs well even at low coverage PacBio read (<10x) providing a cost effective tool to study the dynamics and impact of TEs in natural populations.

## ACKNOWLEDGMENT

The authors wish to thank Nicolas Pollet and Jean-Michel Rossignol for their helpful comments.

